# The role of explicit strategies during reinforcement-based motor learning

**DOI:** 10.1101/234534

**Authors:** Peter Holland, Olivier Codol

## Abstract

Despite increasing interest in the role of reward in motor learning, the underlying mechanisms remain ill-defined. In particular, the relevance of explicit strategies to reward-based motor learning is unclear. To address this, we examined subject’s (n=30) ability to learn to compensate for a gradually introduced 25° visuomotor rotation with only reward-based feedback (binary success/failure). Only two-thirds of subjects (n=20) were successful at the maximum angle. The remaining subjects initially follow the rotation but after a variable number of trials begin to reach at an insufficiently large angle and subsequently return to near baseline performance (n=10). Furthermore, those that were successful accomplished this largely via the use of strategies, evidenced by a large reduction in reach angle when asked to remove any strategy they employed. However, both groups display a small degree of remaining retention even after the removal of strategies. All subjects made greater and more variable changes in reach angle following incorrect (unrewarded) trials. However, subjects who failed to learn showed decreased sensitivity to errors, even in the initial period in which they followed the rotation, a pattern previously found in Parkinsonian patients. In a second experiment, the addition of a secondary mental rotation task completely abolished learning (n=10), whilst a control group replicated the results of the first experiment (n=10). These results emphasize a pivotal role of strategy-use during reinforcement-based motor learning and the susceptibility of this form of learning to disruption has important implications for its potential therapeutic benefits.

## Introduction

The motor system’s ability to adapt to changes in the environment is essential for maintaining accurate movements (Tseng et al., 2007). Such adaptive behavior is thought to involve several distinct learning systems (Haith and Krakauer, 2013; Izawa and Shadmehr, 2011; Smith et al., 2006). For example, the two-state model proposed by Smith et al. (2006) has been able to explain a range of results in force-field adaptation paradigms in which a force is applied to perturb a reaching movement. The model states that learning is accomplished via both ‘fast’ and ‘slow’ processes, the ‘fast’ process learns rapidly but has poor retention, whereas the ‘slow’ process learns more slowly but retains this information over a longer timescale. Subsequently using a visuomotor rotation paradigm, in which the visible direction of a cursor is rotated from the actual direction of hand movement, it has been suggested that the ‘fast’ process resembles explicit re-aiming whereas the ‘slow’ process is implicit (McDougle et al., 2015). The implicit aspect may be composed of several different processes (McDougle et al., 2015), the first and most widely researched being cerebellar adaptation (Izawa et al., 2012). However, additional processes such as use-dependent plasticity and reinforcement of actions that lead to task success are required to fully explain experimental findings (Huang et al., 2014). Haith and Krakauer (2013) have proposed a scheme based on these four processes that attempts a synthesis between the principles of motor learning and the distinction between model-based and model-free mechanisms proposed for reinforcement learning and decision-making (Doll et al., 2016).

The addition of rewarding feedback has proven beneficial in increasing retention of adaptation (Galea et al., 2015; Shmuelof et al., 2012; Therrien et al., 2016) and motor skills (Abe et al., 2011; Dayan et al., 2014). Findings such as these have generated interest in the possibility that the addition of reward to rehabilitation regimes may improve the length of time that adaptations are maintained after training (Quattrocchi et al., 2017; Shmuelof et al., 2012). However, it is still unclear which of the multiple systems mediating motor learning reward may be acting on. Motor learning via purely reward based feedback is also possible and has been applied in two separate forms: binary and graded. Graded point based reward is often based on the distance of the reaching movement from the target and provides information about the magnitude but not the direction of the error (Manley et al., 2014; Nikooyan and Ahmed, 2015). Graded feedback has proved sufficient for learning abrupt rotations (Nikooyan and Ahmed, 2015), however, in certain conditions explicit awareness is required for successful learning (Manley et al., 2014). An alternative method is to only provide binary feedback in which the reward signals task success, such as hitting a target (Izawa and Shadmehr, 2011; Pekny et al., 2015; Therrien et al., 2016). In contrast to graded feedback, only gradually introduced perturbations have successfully been learnt via binary feedback alone (van der Kooij and Overvliet, 2016) and the role of explicit awareness has yet to be examined.

In classical visuomotor adaptation, in which full visual feedback of the cursor is available, gradual adaptation is considered to be largely implicit (Galea et al., 2010). However, this may not be the case when only end-point feedback is provided (Saijo and Gomi, 2010). The question remains as to whether learning a gradually introduced visuomotor rotation based on binary feedback also mainly involves implicit processes. Various methods (Huberdeau et al., 2015) have been used to separate the implicit and explicit components of learning such as asking subjects to verbally report aiming directions (McDougle et al., 2015; Taylor et al., 2014) and forcing subjects to move at reduced reaction times (Haith et al., 2015; Leow et al., 2017). In the current paradigm, we assessed the contribution of strategies at the end of the learning period by removing all feedback but asking subjects to maintain their performance. Subsequently, we asked subjects to remove any strategy they may have been using. Such an approach has previously been used to measure the relative implicit and explicit components of adaptation to different sizes of visuomotor rotations (Werner et al., 2015).

Our second approach to investigating the explicit contribution to learning based on binary feedback was the introduction of a dual task in order to divide cognitive load and suppress the use of strategies. Dual task designs have previously successfully been employed to disrupt explicit processes in adaptation (Galea et al., 2010; Taylor and Thoroughman, 2007, 2008), sequence learning (Brown and Robertson, 2007) and motor skill learning (Liao and Masters, 2001). Various forms of dual task have been used such as counting auditory stimuli (Maxwell et al., 2001), repeating an auditory stimulus (Galea et al., 2010) or recalling words from a memorized list (Keisler and Shadmehr, 2010). We selected a mental rotation task based on using an electronic library of three-dimensional shapes (Peters and Battista, 2008; Shepard and Metzler, 1971). This particular task was selected in order to maximize the likelihood of interfering with the explicit re-aiming process. Indeed, it has previously been shown that both spatial working memory and mental rotation ability correlate with performance in the early ‘fast’ phase of adaptation (Anguera et al., 2009; Christou et al., 2016). Furthermore, the same prefrontal regions are activated during the early phase of adaptation and during the performance of a mental rotation task (Anguera et al., 2009). It has also been suggested that the explicit process of re-aiming in response to visuomotor rotations may involve a mental rotation of the required movement direction (Georgopoulos and Massey, 1987)

If the learning of a gradually introduced rotation via binary feedback is dominated by explicit processes, this should be evidenced by a large change in performance when subjects are asked to remove any strategy. Furthermore, the dual task should severely disrupt learning and could possibly unmask any implicit process.

## Materials and Methods

### Subjects

Sixty healthy volunteers aged between 18 and 35 participated in the study. Forty subjects (thirty-seven females, mean age = 19.9 years) completed experiment 1 and twenty (fifteen females, mean age = 21.6 years) in experiment 2. All subjects were right-handed with no history of neurological or motor impairment and had normal or corrected-normal vision. Volunteers were recruited from the undergraduate pool in the School of Psychology and wider student population at the University of Birmingham and all gave written informed consent. Subjects were remunerated with their choice of either course credits or money (£7.50/hour). The study was approved by the local ethics committee of the University of Birmingham and performed in accordance with those guidelines.

### Experimental Protocol

A similar paradigm has previously been employed and the current protocol was designed to replicate this as closely as possible (Therrien et al., 2016). In addition to the rotation of 15°, we extended this paradigm to a 25° rotation. Subjects performed reaching movements with their right arm using a KINARM (B-KIN Technologies), Figure 1A. Subjects were seated in front of a horizontally placed mirror that reflected the visual stimuli presented on a screen above (60 Hz refresh rate). Reaching movements were performed in the horizontal plane whilst subjects held the handle of a robotic manipulandum, with the arm hidden from view by the mirror.

**Figure 1.**
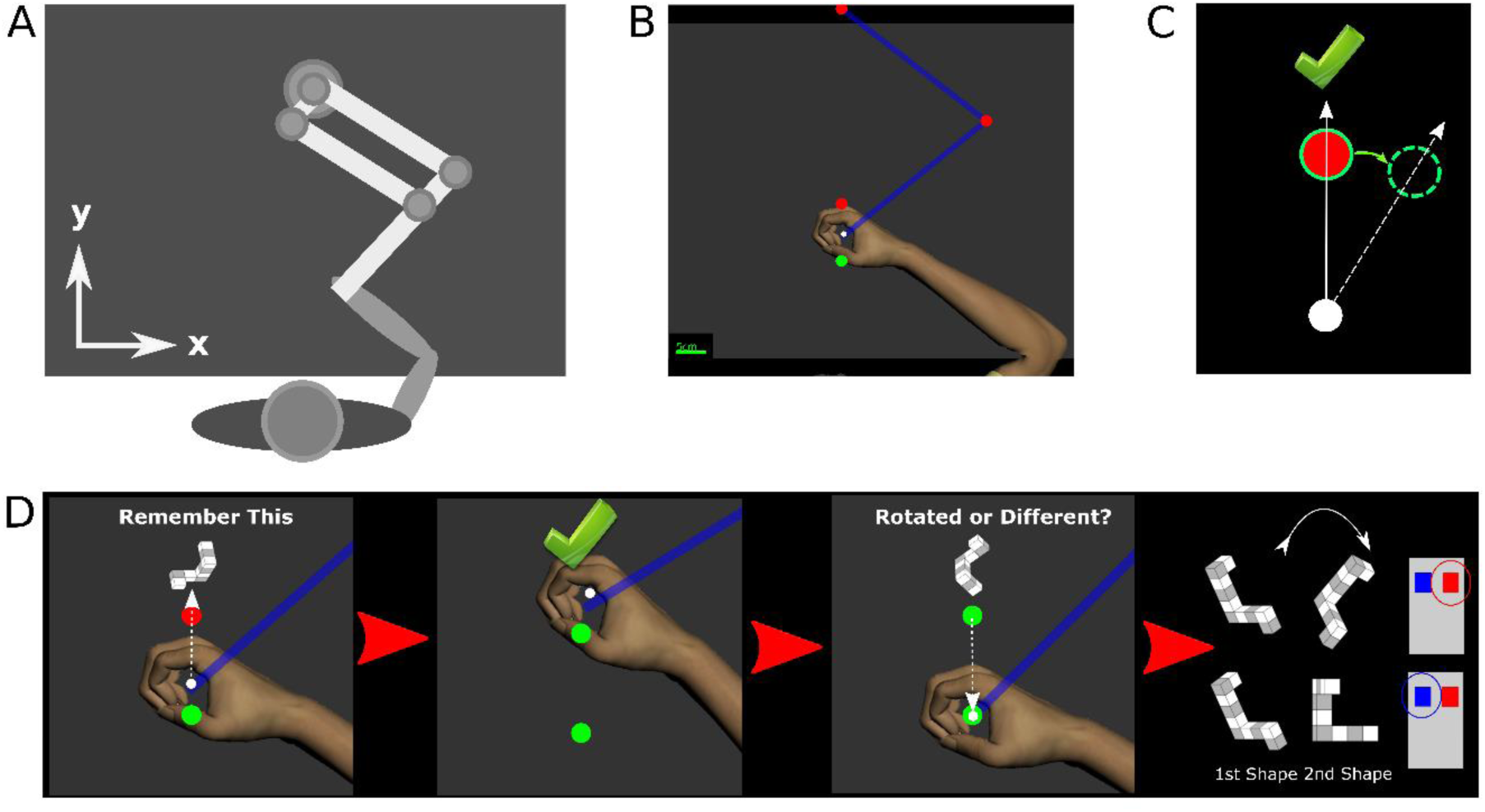
*Experimental design.* ***A**, Subjects held the handle of robotic manipulandum with their right hand, the position of the arm and handle was hidden from sight and feedback was provided on a horizontal screen. **B**, Subjects made ‘shooting’ movements from a starting position (green circle) towards a target (red circle), after the initial practice trials the position of the cursor (white circle) was no longer visible at any point. **C,** Successful trials were indicated to the subject with the display of a green tick after the cursor had passed through a region centered on the target, over the course of the paradigm the position of the reward region gradually moved (solid green circle to dashed green circle) whilst the visible target (red circle) remained in the central location. By the end of the learning period a successful reach (dotted white line) was rotated by a maximum of either 15° or 25 °. **D,** Time-course of Experiment 2, at the same time as the target appeared on screen a ‘shape’ was also displayed slightly above it, the subject was asked to memorize this shape. After the reach was completed and the hand returned to the starting position subjects used their left hand to respond with a button press as to whether they believed the new shape shown on screen was a rotated version of the shape or an entirely different shape.*

### Experiment 1

Two different paradigms were employed in Experiment 1, both consisted of a gradually introduced rotation of the required angle of reach for a trial to be considered successful. The maximal extent of the rotation was either 15° (n=10) or 25° (n=30). Subjects were required to learn the rotation on the basis of only binary feedback indicating if they had successfully hit the target region. After the rotation had reached the maximal extent, all feedback was extinguished and two further blocks of trials were performed to assay the level of retention and to what extent this was explicit in nature.

A total of 670 or 470 trials were performed for the 25° and 15° paradigms, respectively. Each trial followed an identical sequence. Initially a starting position was displayed on screen (red colored circle, 1cm radius), after subjects had moved the position of the cursor (white circle, 0.5cm radius) into the starting position, the starting position changed color from red to green. After a small delay (randomly generated, 500-700ms), in which subjects had to maintain the position of the cursor within the starting circle, a target (red circle, 1cm radius) appeared directly in front of the starting circle at a distance of 10cm. Subjects were instructed to make rapid ‘shooting’ movements that intercepted a visual target, they were instructed that they did not have to attempt to terminate their movement in the target but pass directly through it (Figure 1B). If the cursor intercepted a ‘reward region’ (±5.67°), initially centered on the visible target, the movement was considered successful and the target changed color from red to green and a large (8x8cm) green ‘tick’ was displayed at a distance of 20cm directly in front of the starting position (Figure 1C). However, if the cursor did not intercept the reward region the trial was considered unsuccessful and the visible target disappeared from view. Movement times, defined as the time from leaving the starting circle to reaching a radial distance of 10cm, were constrained to a range of 200-1000ms. Movements outside of this range but at the correct angle were counted as incorrect trials and no tick was displayed. As a visual cue, movements outside of the acceptable duration were signaled with a change of the target color, blue for too slow and yellow for too fast. After the completion of a reaching movement the robot returned the handle to the start position and subjects were instructed to passively allow this whilst maintaining their grip on the handle. Reaction times, defined as the difference in time between the appearance of the target and the time at which the cursor left the starting circle, were limited to a maximum 600ms. If a movement was not initiated before this time the target disappeared and the next trial began after a small delay and these trials were excluded from further analysis.

After an initial period of ten trials, in which the cursor position was constantly visible, for the remainder of the experiment it was extinguished. The only feedback subjects received was a binary (success/fail) signal indicating if the angle of reach was correct, in the form of a change of target color and the appearance of the tick. For an initial period of forty trials the reward region remained centered on the position of the visual target, after this it was shifted in steps of 1° every twenty trials. This manipulation ensured that for a reaching movement to be considered correct it must be made at an increasingly rotated angle from the visual target (Figure 1C). Subjects were pseudo-randomly assigned to groups that received either a clockwise or counter-clockwise rotation. Once the reward region had reached the maximal angle, either 15° or 25°, it was held constant for an additional twenty trials. Subsequently, subjects were informed that they would no longer receive any feedback about their performance but that they should continue to perform in the same manner as before, this ‘Maintain’ block consisted of fifty trials. Following this, subjects were asked a series of simple questions to assay their awareness of the rotation, answers were noted by the experimenter. Subsequently all subjects were told ‘During the task we secretly moved the position of the target that you had to hit. You will still not receive information on whether you hit the target or not but please try to move as you did at the start of the experiment’. Crucially subjects were not informed of the direction or magnitude of the rotation they had experienced. The final ‘Remove’ block consisted of fifty trials. The position of the handle throughout the task was recorded at a sampling rate of 1 kHz and saved for offline analysis.

### Experiment 2

Experiment 2 comprised of the same reaching task as Experiment 1 but with the addition of a mental rotation dual task. The dual task required subjects to hold a three-dimensional shape in working memory for the duration of the reaching movement (Figure 1D). Subjects had to respond with a button press using their left hand to indicate if a shape displayed at the end of the reaching movement was a rotated version of a shape displayed at the time of target presentation or a different shape.

Shapes had the form of a series of connected cubes, alternately colored grey and white, they were selected from an electronic library designed on the basis of the Shepard and Metzler type stimuli (Peters and Battista, 2008; Shepard and Metzler, 1971). All rotations were performed within the plane of the screen, i.e. although the stimuli represented three-dimensional shapes all rotations were in two-dimensions. A subset of 26 shapes were selected from the library for use in this experiment and are available on https://osf.io/vwr7c/. The trial protocol was the same as that employed in Experiment 1 but at the time when the target circle appeared, a randomly selected shape from the subset was displayed in an 8x8cm region at a position 20cm away from the starting position. Subjects were instructed to commit this shape to memory. The shape remained visible on screen until the end of the reaching movement, the point at which the radial amplitude of the cursor exceeded 10cm. The shape was then extinguished and the same binary feedback as employed in Experiment 1 was displayed. After the robot had guided the handle back to the starting position a second shape was displayed. In half of the trials this was an identical shape to the first one but had undergone a rotation selected at random from a uniform distribution of 0-360°, in the other half of trials it was a different shape selected at random from the library. The order of trials in which the shape was either rotated or different was randomized and subjects had a maximum of 2s to respond. Subjects in the Dual Task group (n=10) were instructed to press the right-sided button of two buttons on a button box held in their left hand if they believed the second shape to be a rotated version of the first one and the leftsided button if they believed it was a different shape. Importantly subjects were given no feedback on their performance in the dual task but were informed prior to the experiment that this would be monitored, the responses were recorded and analyzed offline. This design was selected in order to avoid any interfering effects of rewarding feedback from the dual task with the binary feedback in the reaching task. As a control, another group of subjects received identical visual stimuli but were instructed to press a random button of the two on each trial. Subjects were pseudo-randomly assigned to either the Control or Dual Task groups.

For Experiment 2 the familiarization period at the start of the experiment, in which the position of the cursor was visible, was extended to twenty trials in order for subjects to have sufficient time to acclimatize to the additional timing requirements of the button press. The paradigm subsequently followed that of Experiment 1 with a maximal angle of rotation of 25°.

### Data Analysis

All data analysis was performed with custom written routines in MATLAB (The Mathworks) and extracted data and all code required to reproduce the analysis and figures in this paper are freely available on (https://osf.io/vwr7c/).

The end point angle of each reaching movement was calculated either at the time that the cursor intercepted the reward region or in the case of incorrect trials when the cursor reached a radial amplitude of 10cm. An angle of zero degrees was defined as a movement directly ahead, i.e. toward the visible target position. A positive angle of rotation was defined as a clockwise shift of the reward region, and reach angles and target positions for the counter-clockwise rotation were sign-transformed to positive values for comparability. The ‘Baseline’ period was defined as the first forty trials without visual feedback of the cursor, during which the reward region was centered on the visual target. Subjects were considered to have successfully learnt the rotation if the mean end point angle of the reaching movements fell within the reward region during the last twenty trials before the ‘Maintain’ period, a time at which the rotation was held constant at its maximal value.

During the retention phase of the experiment (last one hundred trials), we calculated the amount of retention that could be accounted for by explicit and implicit processes. A subject’s implicit retention was defined as the difference between the mean reach angle in the final fifty trials (‘Remove’ blocks), after subjects had been instructed to remove any strategy they had been using, and mean reach angle during the ‘Baseline’ blocks. A subject’s explicit retention was defined as the difference between the mean reach angle during the ‘Maintain’ blocks, the first fifty trials after removal of binary feedback in which subjects were instructed to continue reaching as before, and the implicit retention.

In order to analyze the effect of reward on subjects behavior we conducted trial-by-trial analysis in a manner similar to one that has previously been employed for analysis of reaching performance in response to binary feedback (Pekny et al., 2015). The change in reach angle following trial n, Δ*u*^(*n*)^, was defined as the difference between consecutive trials:

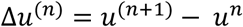

Subsequently we examined the distributions of *Δu* following only rewarded (correct) or unrewarded (wrong) trials. The resulting distributions of *Δu* were non-normal and therefore we analyzed and report the median and median absolute deviation (MAD) of each subject’s distributions. We also examined the absolute change in reach angle |*Δu*|, i.e. the magnitude of change regardless of direction.

In order to investigate the effects of a reward history spanning multiple trials we examined the |*Δu*| following all possible combinations of success in the previous three trials. We first searched each subject’s responses for the occurrence of all eight possible sequences of reward and calculated the mean change in reach angle following each. We then quantified this behavior using a state-space model in which |*Δu*| was a function of the outcome of the previous three trials as well as variability (*ε*) that could not be accounted for by the recent outcomes (Pekny et al., 2015):

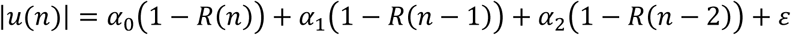

In the above equation *R* represents the presence of reward on a given trial with a value of 1 for a correct trial, *R*(*n*) therefore represents the presence of reward on the previous trial with *R(n −* 1) and *R*(*n* − 2) the preceding two trials. The components *α*_0_, *α_1_* and *α_2_* represent the sensitivity to the outcomes of these trials with higher values indicating subjects made larger changes in response to the outcome of that trial.

The verbal responses to the questions asked before the start of the ‘Remove’ block was noted down by the experimenter and analyzed offline. A subject’s awareness of the perturbation and efforts to deliberately counter it were rated on a scale of 0, 0.5 and 1, with 0 indicating no awareness and 1 full awareness.

### Statistical Analysis

Statistical analysis was performed in MATLAB. In order to test for initial effects mixed design ANOVAs were used, with Group (25RotSucces, 25RotFail etc.) as the between-subjects factor and time-point (Baseline, 15° Block, Maintain etc.) or MeasuredVariable (Median *Δu*, Reward Component etc.) as the within-subjects factor. The Greenhouse-Geiser correction was applied in cases of violation of sphericity and corrected p-values and degrees of freedom are reported in the text. In cases in which a significant interaction was found in the ANOVA, post-hoc tests were performed to test for differences between groups at each TimePoint or MeasuredVariable. As data was often found to be non-normally distributed using Kolmogorov-Smirnov tests, the non-parametric Kruskal-Wallis test was applied throughout. In cases of a significant effect of group on an individual outcome measure, further pairwise comparisons of mean group ranks were employed and Bonferroni corrected p-values are reported in the text. For tests of a difference of a single group from zero, such as in testing for implicit learning, Wilcoxon-Signed Rank tests were employed and Bonferroni corrected p-values are reported in the text. A critical significance level of α=0.05 was used to determine statistical significance.

## Results

### Experiment 1: Successfully learning to compensate for a 25° rotation includes a large explicit component

We first sought to investigate the size of a gradual introduced visuomotor rotation that subjects can learn based on binary feedback. All subjects who experienced the 15° rotation (15Rot group) learnt to fully compensate (Figure 2A). Successful compensation was defined as having a mean reach angle within the reward region in the final twenty trials before the retention phase. However, for the 25° group (25Rot, magenta group, Figure 2B), the average reach direction fell outside the reward region, indicating incomplete learning. Underlying the mean performance was a split in behavior: some subjects successfully learnt the full rotation, whereas one third of subjects did not. On the basis of this behavior, they were categorized into two subgroups: 25RotSuccess (red group, N=20) and 25RotFail (blue group, N=10), respectively.

**Figure 2.**
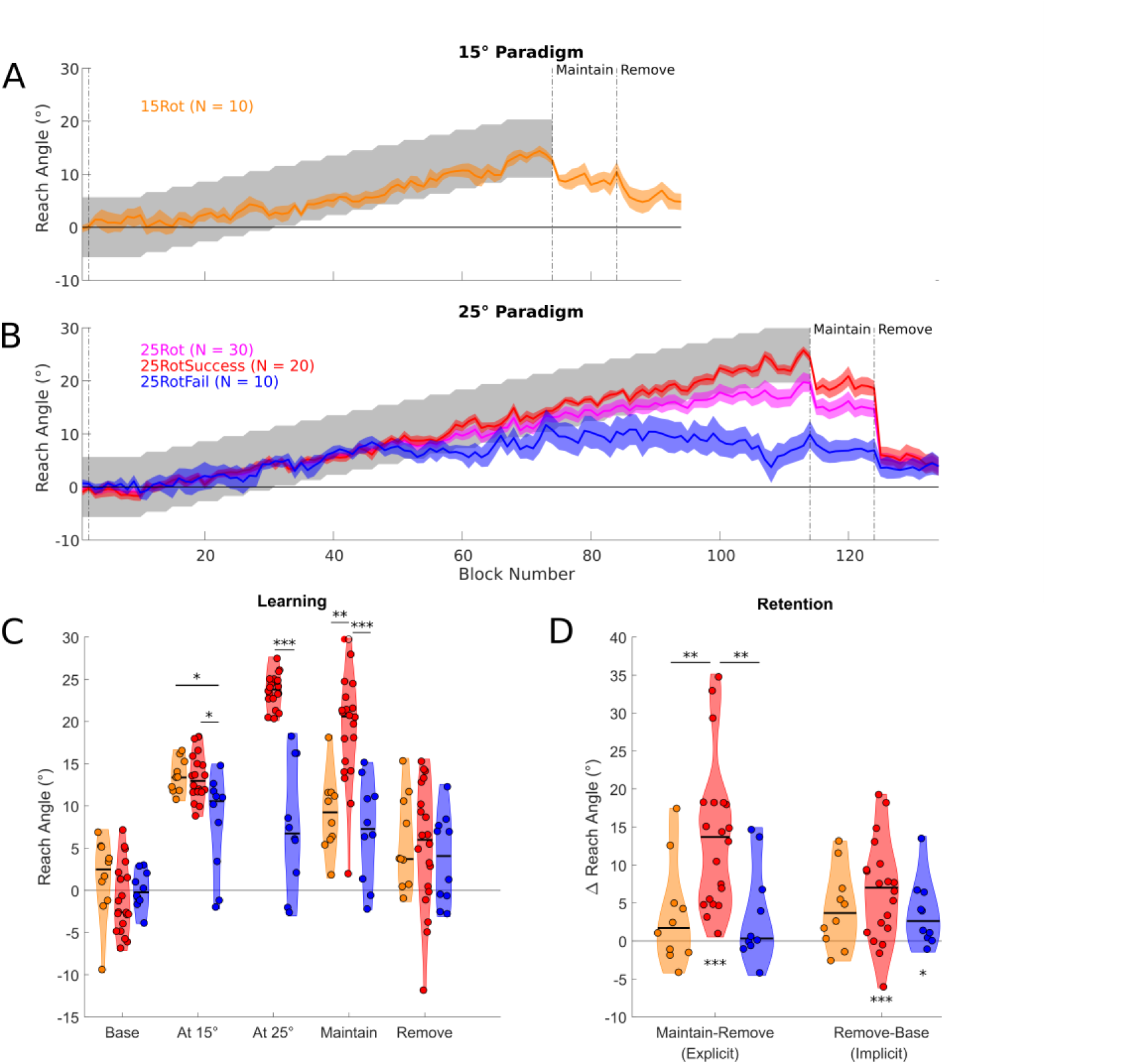
*Experiment 1: group performance.* ***A,** Reach angle averaged over blocks of 5 trials, solid colored lines represent the mean of each group and the shaded region represents SEM. The average behavior of subjects in the 15Rot paradigm (Orange) fell consistently within the rewarded region (grey shaded region) indicating successful learning. **B**, Average reach angle over blocks for all subjects in the 25Rot paradigm (magenta) and also the same subjects split into two groups based on success at the final angle (25RotSuccess – red, 25RotFail – blue). **C,** Distribution plots displaying the reach angles for subjects in the three groups at various timepoints throughout the experiment with individual data points overlaid on an estimate of the distribution. Horizontal black line in the distribution represents the group median. **D,** Distribution plots of the computed variables of Implicit (‘Remove-Baseline’) and Explicit (‘Maintain-Implicit’) retention. Significance stars above horizontal black bars indicate differences between the groups (* P < 0.05, ** *P < 0.01,* *** *P* <0.001). Significance stars below the distributions represent a significant difference from zero.*

Next, we compared reach angle for the three groups (15Rot, 25RotSuccess and 25RotFail) at specific time points in order to gain an understanding at which stage the difference emerged (Figure 2C, D). Despite no difference between groups at baseline (H(2) = 4.03, p = 0.13, Kruskal Wallis), a difference had emerged at 15 degrees (H(2) = 9.63, p = 0.008; Figure 2C). Specifically, reach angle for the 25RotFail group was lower than both the 15Rot (p = 0.022) and the 25RotSuccess groups (p = 0.014). During the ‘Maintain’ phase, when binary feedback had been removed but subjects were instructed to continue reaching as before, there was a significant effect of group (H(2) = 20.08, p < 0.001; Figure 2B, C). Unsurprisingly, the 25RotSuccess group was greater than the 15Rot (p = 0.002) and the 25RotFail groups (p < 0.001). Crucially, after subjects were instructed to remove any strategy and reach as they did at the beginning of the experiment, there was no difference between the groups (H(2) = 0.78, p = 0.68; Figure 2B, C). Analysis of the reach angles during the paradigm revealed that even at a rotation of 15° there was divergence between the 25RotFail and 25RotSuccess groups. Furthermore, the instruction to remove any strategy resulted in a return to a similar level of performance across all three groups.

We probed the nature of learning by calculating the implicit and explicit components of retention (Figure 2D). Implicit retention reflected the retention after removal of any strategies, whereas Explicit retention represented the change in behavior accounted for by the use of strategies. The Explicit component of the 25RotSuccess group was greater than both 15Rot (p = 0.006) and 25RotFail (p = 0.006). Furthermore, only the 25RotSuccess (Z = 210, p < 0.001) group had a significant Explicit component to their retention. Whilst there was no effect of Group on the Implicit component (H(2) = 1.84, p = 0.40), both groups in the 25° paradigm showed a significant difference from 0 (25RotSuccess, Z = 193, p = 0.001; 25RotFail, Z = 48, p = 0.014), however, the 15Rot group was no longer significant after correction for multiple comparisons (Z = 48, uncorrected p = 0.037, corrected p = 0.111). Therefore, whilst all three groups showed a similar small level of implicit retention, only the subjects who successfully learnt the 25° rotation showed evidence for explicit learning.

In order to understand the mechanism of learning, and how this might differ between the 25RotSuccess and 25RotFail groups, we examined trial-by-trial behavior. Two distinct types of behavior were apparent (Figure 3). Behavior in those that failed (Figure 3B) was initially similar to successful subjects (Figure 3A) but at some point subjects began to fail to reach at a sufficient angle. Subsequently the angle of reach returned to near zero, despite a continued lack of reward. The angles at which subjects in the 25RotFail group failed varied (mean=13.0°), but all displayed the same pattern of return to baseline (Figure 3C). Given the apparently similar behavior in the initial learning stage, it is important to know whether there are differences even at this early stage. To this end, we only included trials in the initial successful period for the 25RotFail group in all subsequent analysis of trial-by-trial behavior, i.e. trials on the left-hand side of the vertical colored line for each subject (Figure 3C). For the 25RotSuccess and 15Rot groups all trials during the learning period were analyzed. Crucially, there was no difference in the percentage of correct trials within this period between the groups (H(2) = 2.19, p = 0.33).

**Figure 3.**
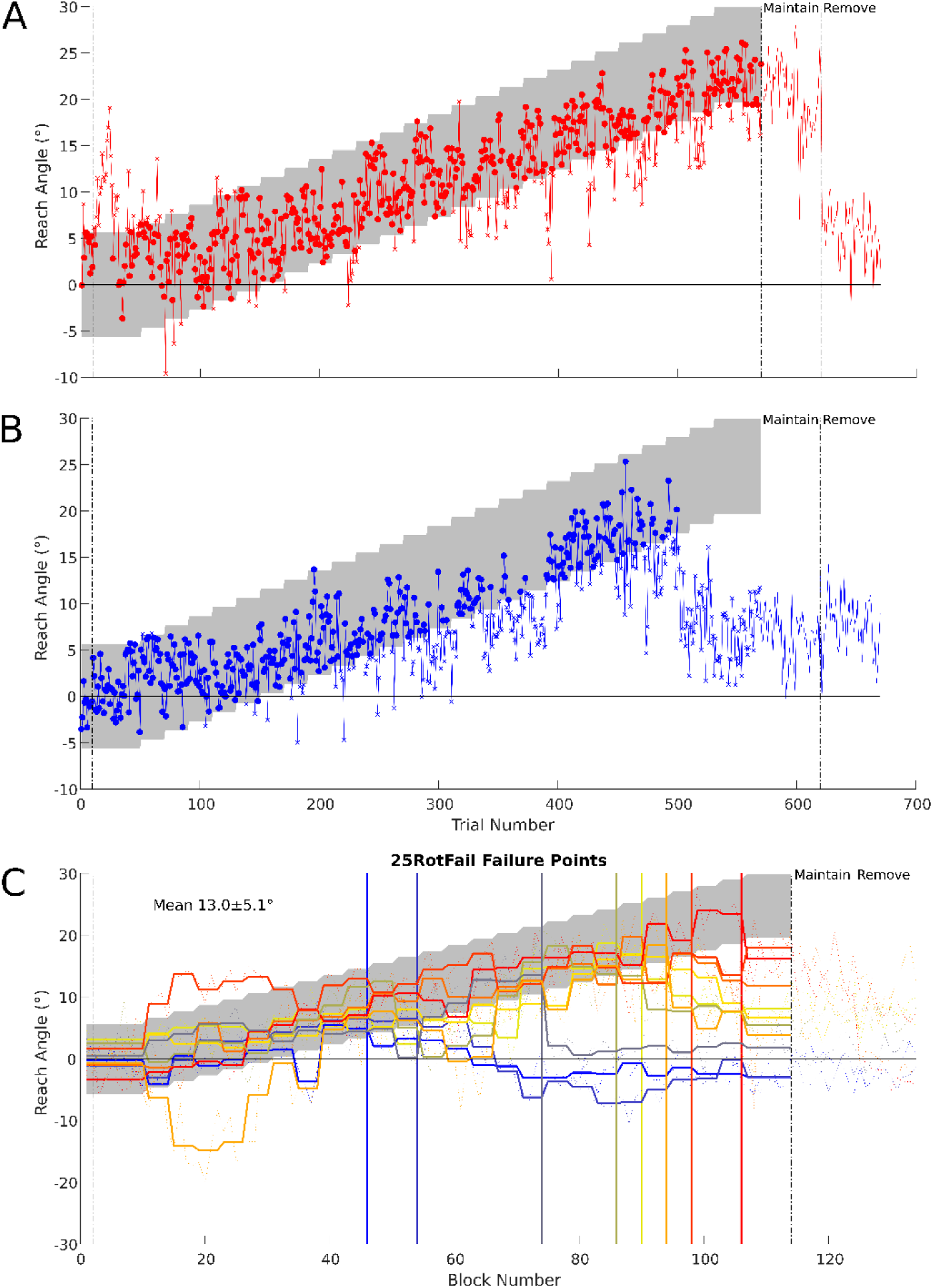
*Experiment 1: trial-by-trial behavior.* *Example of trial by trial reach angles from a subject who was successful at the final angle (**A**) and one who was unsuccessful (**B**). In each case rewarded trials are indicated with a circular marker and non-rewarded trials with a ‘x’. The grey shaded region indicates the reward region. **C,** Failure points for subjects in the 25RotFail group, thick lines are the mean reach angle for each subject at each rotation angle, thin lines represent mean of each block (average of 5 trials), colors go from hot to cold matching failure angles ranging from high to low. Vertical lines represent the last angle at which mean reach fell within rewarded region for each subject. The mean and standard deviation of all angles of failure is displayed as text.*

Next, we examined if changes in reach angle were affected by the outcome of the previous trial. A similar analysis has been employed previously (Pekny et al., 2015). We examined the distributions of *Δu* following only rewarded (Correct) or unrewarded (Wrong) trials. The resulting distributions of *Δu* were non-normal and therefore we report the median and median absolute deviation from the median (MAD). Whilst the median *Δu* was greater following unrewarded trials (F(1,37) = 119.80, p < 0.001; Figure 4A), this effect was similar across groups (F(2,37) = 1.18, p = 0.64). Similarly, the MAD of *Δu* was also greater following Wrong trials, indicating that not only did all groups make larger changes in reach angle but also that there was greater variability in these changes (Figure 4B). Despite a significant interaction with Group (F(2,37) = 5.32, p = 0.019), the trend for a higher MAD of *Δu* following Wrong trials for the 25RotSuccess group (Figure 4B) did not reach significance after correction for multiple comparisons (H(2) = 5.63, p = 0.06). Subsequently we repeated the analysis but considered the absolute change in reach angle (|*Δu*|, Figure 4C, D). Here there was a significant interaction with Group for both median |*Δu*| (F(2,37) = 7.89, p = 0.003) and MAD of |*Δu*| (F(2,37) = 7.39, p = 0.004) following Wrong trials. Post-hoc tests revealed that the 25RotSuccess group displayed a significantly greater median |*Δu*| (p = 0.024) and MAD of |*Δu*| (p = 0.035) than the 25RotFail group. There was no difference between the groups in the magnitude or variability of the change in reach angle after correct trials. The analysis of the absolute changes in reach angle reveal that even during the period in which they are successful, the 25RotFail group made smaller and less variable changes following unrewarded trials.

**Figure 4.**
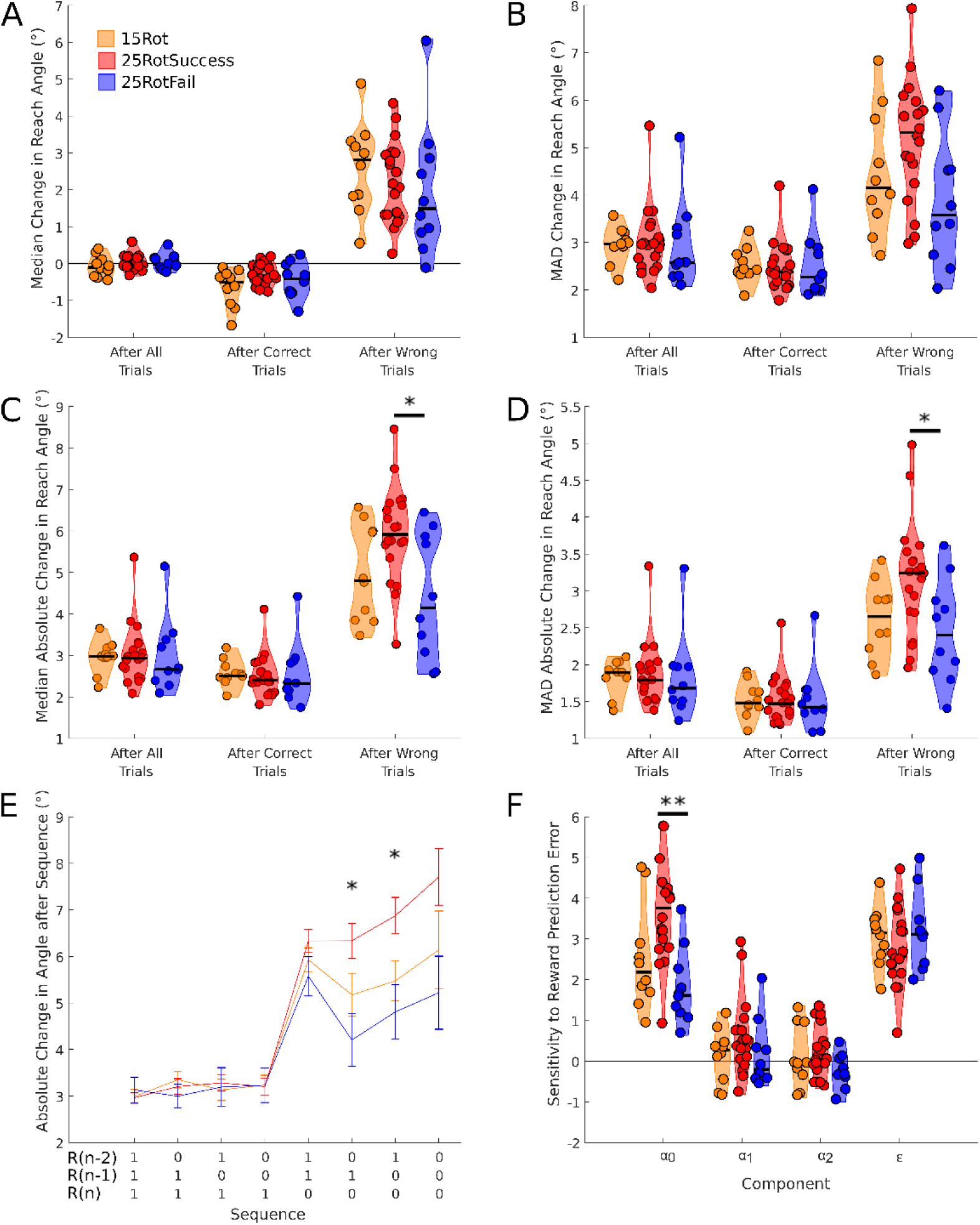
*Experiment 1: performance after correct and incorrect trials.* *Analysis of the effects of the success of the previous trial and reward history on trial by trial changes in reach angle for the three groups in Experiment 1 (15Rot – Orange, 25RotSuccess – Red, 25RotFail – Blue). Median (**A**) and MAD (**B**) of change in reach angle separated by the success of the previous trial. Median (**C**) and MAD (**D**) of the absolute change in reach angle separated by the success of the previous trial. E, The absolute change in reach angle following all combinations of trial success over the previous three trials. F, Sensitivity to the outcomes of each of the previous trials. Significance stars above horizontal black bars indicate differences between the groups* (* *P < 0.05,* ** *P < 0.01).*

In addition to the effect of the previous trial it is possible that subjects are sensitive to a history of outcomes spanning multiple previous trials (Pekny et al., 2015). In order to investigate the effects of reward history we examined the |*Δu*| following all possible combinations of success in the previous three trials (Figure 4E). We quantified this behavior using a state-space model in which |*Δu*| was a function of the outcome of the previous three trials. The components *α_0_*, *α_1_* and *α_2_* represent the sensitivity to the outcome of the last three trials with *α_0_* being the most recent (Figure 4F), *ε* represents variability that could not be accounted for by the recent outcomes. There was an interaction between component and group (F(3.49,64.51) = 4.49, p = 0.004). All groups were most sensitive to the most recent trial outcome (*α_0_*) with the 25RotSuccess group displaying significantly greater change than 25RotFail (p = 0.001). There was no difference between groups for other components indicating that differences in behavior were driven by the sensitivity to the outcome of the most recent trial. From these results it becomes apparent that, even in the initial period of success, subjects who will go on to fail to learn the full rotation show a decreased sensitivity to errors.

There was no difference between groups for either movement time (H(2) = 4.95, p = 0.084) or reaction time (H(2) = 2.98, p = 0.23). Additionally, within the 25RotFail group reaction and movement times did not differ before and after the point of failure (Z = 25, p = 0.85 and Z = 42, p = 0.16 respectively). In response to the questions asked to probe awareness we found no significant difference between the groups (χ^2^(2) = 3.75, p = 0.15).

### Experiment 2: Addition of a dual task prevents learning

Following the finding of Experiment 1 that successful reinforcement-based motor learning involves the development of an explicit strategy, we sought to investigate if it was possible to disrupt learning by dividing cognitive load. To this end, we required subjects to hold a shape in memory during the period of movement (Figure 1D).

The DualTask (N=10) group displayed little learning and none successfully compensated for the maximum rotation (Green group, Figure 5A). As in Experiment 1, the Control (N=10) group on average fell short of complete learning (Purple group, Figure 5A, B), indicated by the mean reach direction falling outside the reward region in the final learning blocks. However, the average of the group obscures a similar split in behavior with only six subjects successfully learning the full rotation and four failing to do so, which we will label (ControlSuccess and ControlFail, respectively (Figure 5B).

**Figure 5.**
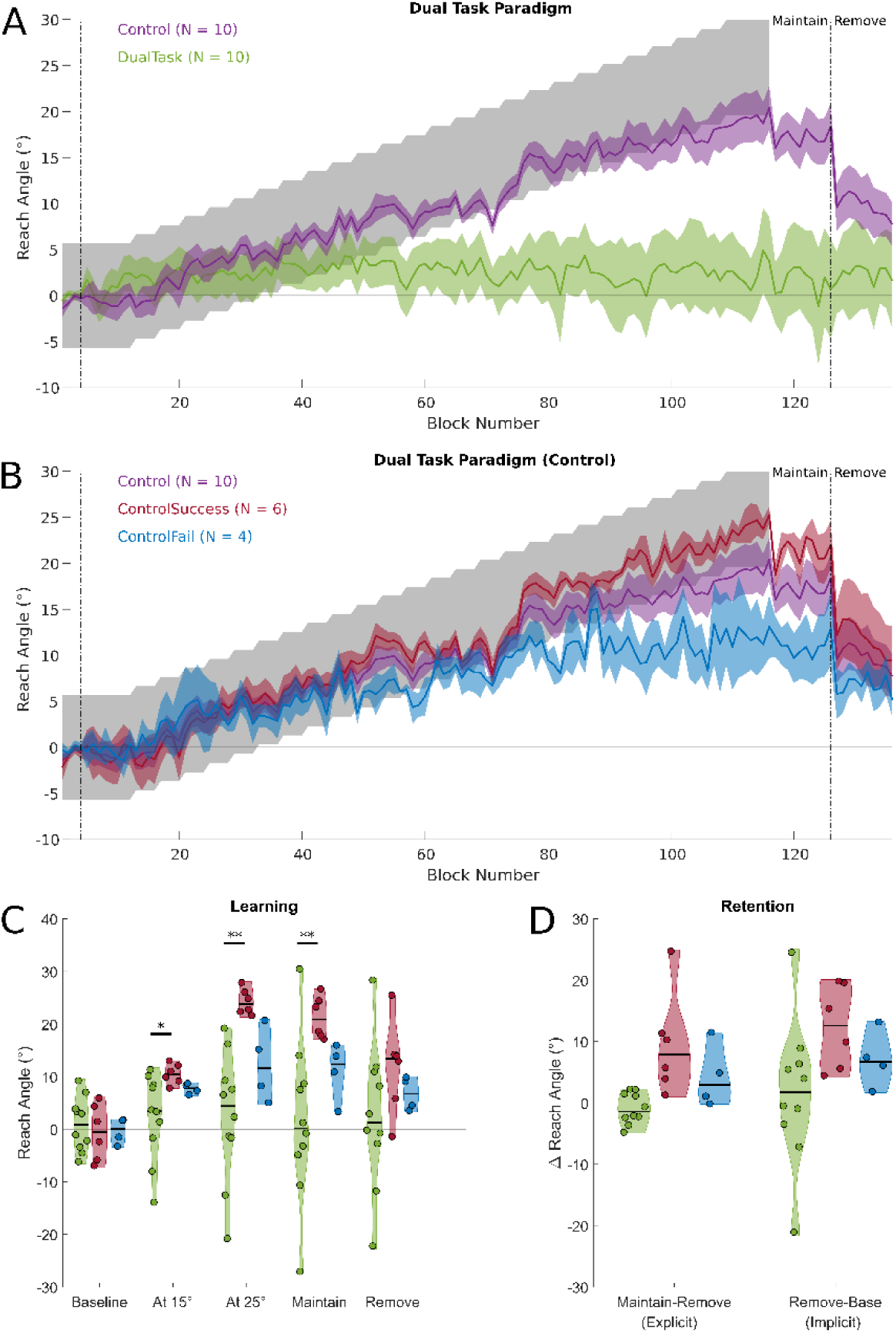
*Experiment 2: group performance.* *Change in reach angle over blocks (average of 5 trials) during the dual task experiment. **A,** Group performance for the DualTask (Green) and Control (Purple) task groups, the line indicates the mean and shaded region the SEM. The grey shaded region represents the reward region. **B,** the split of the control task group into ControlSuccess (Dark Red) and ControlFail (Blue). **C,** Distribution plots displaying the performance at different time points for the dual task, and split control groups. The shaded region represents an estimation of the distribution and is overlaid with data for each individual subject. **D,** Distribution plots of the difference in reach angle during retention phases indicating the implicit and explicit components of retention. Significance stars above horizontal black bars indicate differences between the groups* (* *P < 0.05, **P < 0.01).*

Examining performance in the same time periods as Experiment 1 (Figure 5C) revealed no difference between the three groups at baseline (H(2) = 0.38, p = 0.83). However, by the time the angle of rotation had increased to 15° a significant difference had already emerged (H(2) = 6.88, p = 0.03), with the DualTask group displaying lower reach angle than ControlSuccess (p = 0.011).

As can be seen from the performance of individuals in the DualTask group (Figure 6), there were very few correct trials (mean angle of failure 6.0°) rendering the analysis of trials within the successful period employed for Experiment 1 invalid. Despite this limitation for the DualTask group, the analysis could still elucidate differences between the ControlSuccess and ControlFail groups and reassuringly the mean angle of failure in ControlFail group is 13°, similar to experiment 1. However, the small group numbers preclude statistical comparison between the ControlSuccess and ControlFail groups but the pattern of behavior was visually similar to that in Experiment 1 (Figure 7). Overall the analysis of sensitivity to reward history produced remarkably similar results to Experiment 1 with the primary difference between those who learn and those who fail to do so being the sensitivity to the outcome of the most recent trial (Figure 7F).

**Figure 6.**
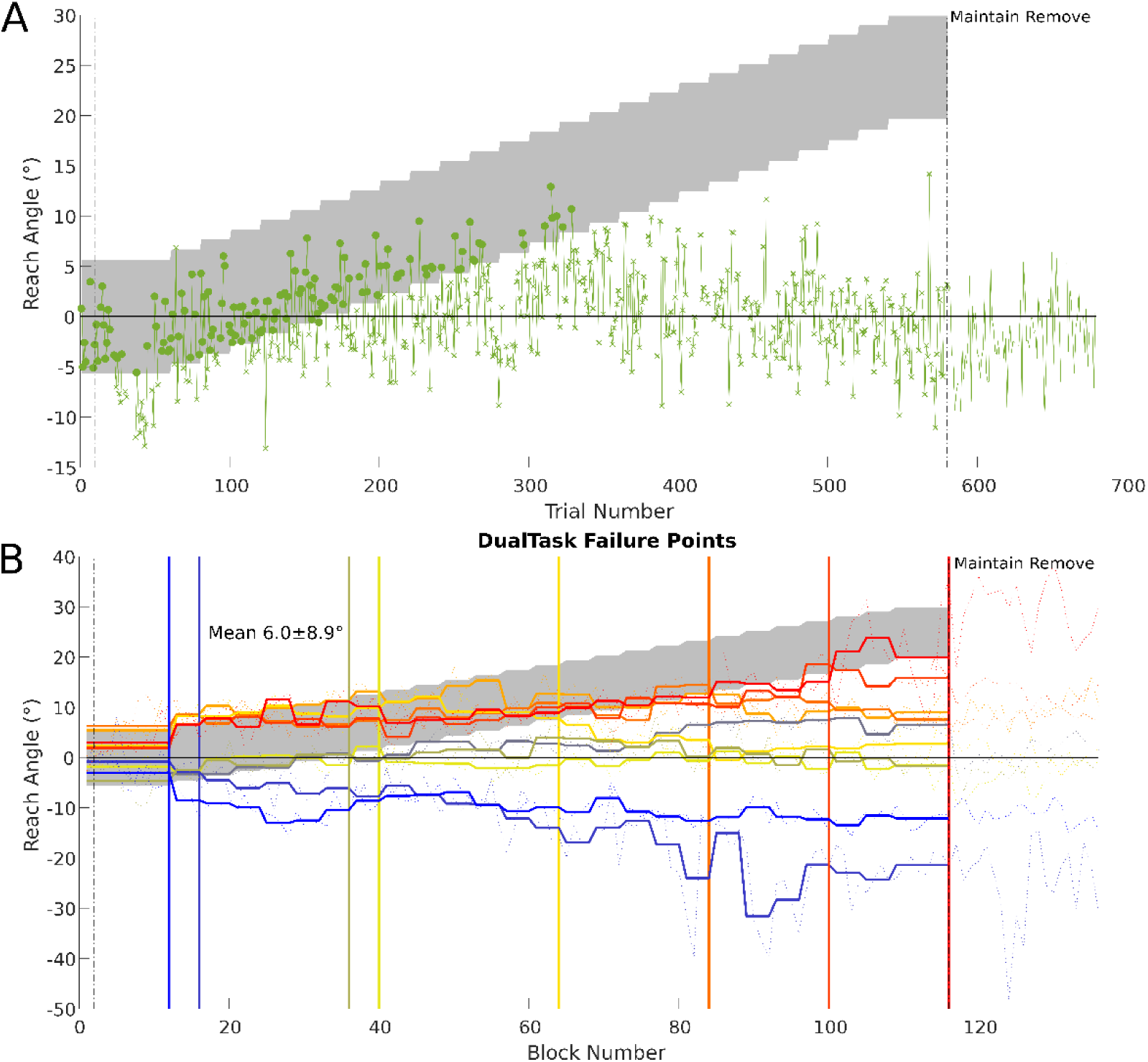
*Experiment 2: trial-by-trial behavior.* *Example of trial by trial reach angles from a subject performing the dual task (**A**) rewarded trials are indicated with a circular marker and non-rewarded trials with a ‘x’. The grey shaded region represents the reward region. **B,** Failure points for subjects in the DualTask group, thick lines are the mean reach angle for each subject at each rotation angle, thin lines represent mean of each block, colors go from hot to cold matching failure angles ranging from high to low. Vertical lines represent the last angle at which mean reach fell within rewarded region for each subject. Th mean and standard deviation of the angle of failure is reported as text in the figure.*

**Figure 7.**
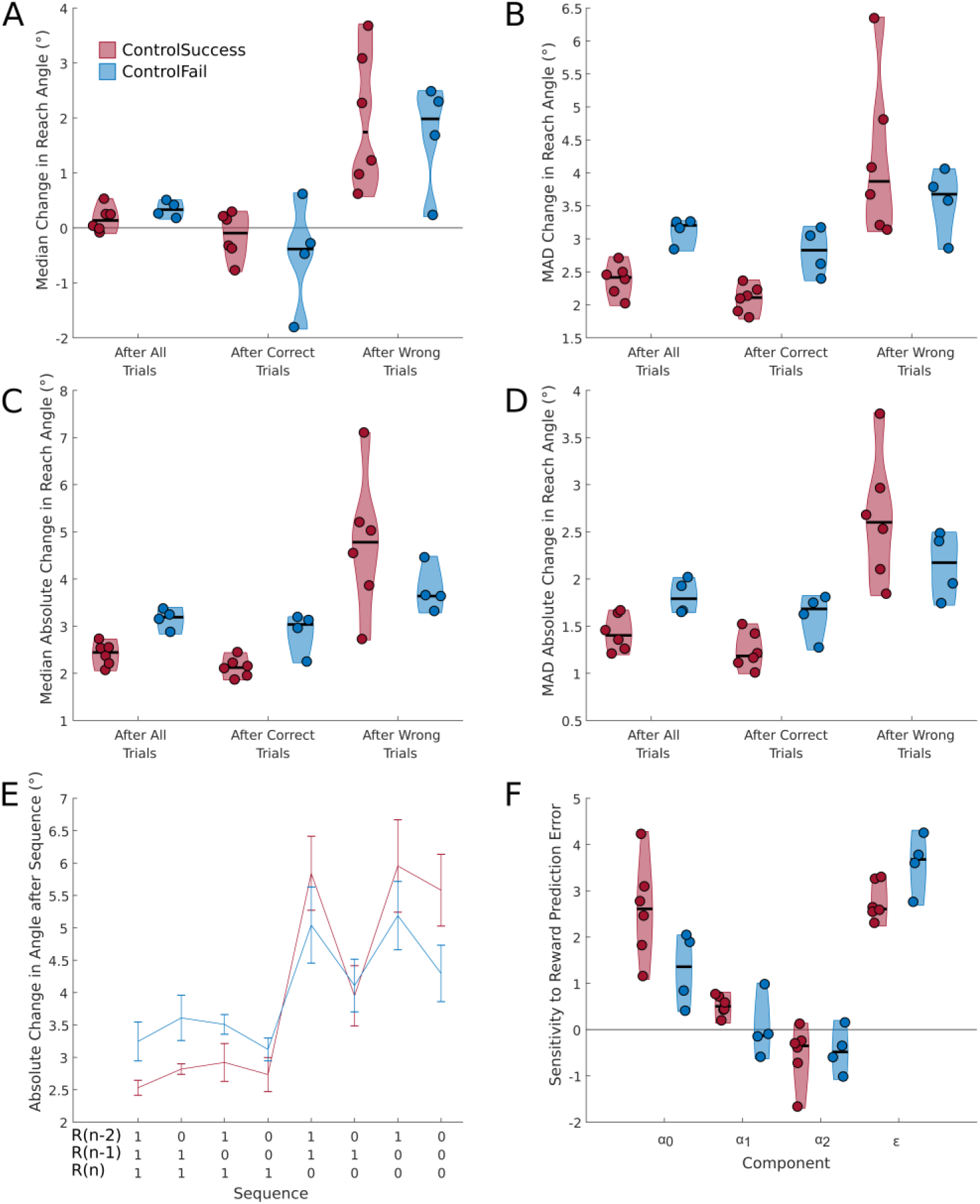
*Experiment 2: performance after correct and incorrect trials.* *Analysis of the effects of the success of the previous trial and reward history on trial by trial changes in reach angle for the two groups performing the control task in Experiment 2. Distribution plots for median (**A**) and MAD (**B**) of change in reach angle separated by the success of the previous trial. Median (**C**) and MAD (**D**) of the absolute change in reach angle separated by the success of the previous trial. **E,** the absolute change in reach angle following all combinations of trial success over the previous three trials. **F,** sensitivity to the outcomes of each of the previous trials.*

Finally, the DualTask subjects successfully engaged in the task mental rotation task as evidenced by a significant difference in percentage of correct button presses (H(2) = 15.30, p < 0.001), the DualTask group responded correctly (67.21 ± 3.60%) more in comparison to the ControlSuccess (p = 0.014) and the ControlFail (p = 0.002) groups. Engagement in the DualTask increased reaction time when compared to ControlSuccess (p = 0.008). There was no effect of Group on movement time (H(2) = 0.64, p = 0.73).

## Discussion

The role of explicit strategies during reinforcement-based motor learning has previously been ill-defined. Here, we reveal that successfully learning to compensate for large, gradually introduced, rotations based on binary (reinforcement-based) feedback involves the development of an explicit strategy, and that not all subjects are able to do so. In both Experiment 1 and the Control group of Experiment 2 only two thirds of subjects were able to successfully learn a large perturbation, and those that did accomplished this principally via the use of a strategy. Analysis of the trial-by-trial behavior indicated that subjects adjusted their motor commands mainly in response to incorrect trials, and that they were most sensitive to errors made in the most recent trial. Subjects who would go on to fail to learn the full rotation exhibited reduced sensitivity to errors, even in the initial period in which they successfully followed the rotation. Further evidence for the explicit nature of the learning in this task was provided by Experiment 2, where increasing cognitive load via the addition of a dual task prevented learning.

Previous experiments investigating the learning of rotations based on binary feedback have employed relatively small angles (Izawa and Shadmehr, 2011; Pekny et al., 2015; Therrien et al., 2016), with the 15° rotation used by Therrien et al. (2016) the largest reported to date. Indeed, when a rotation of 15° was used in Experiment 1 all subjects were successful in fully compensating for the visual rotation. Furthermore, there was no evidence for an explicit component to retention in the subjects who learnt the 15° rotation. In contrast, successful subjects in both experiments with a 25° rotation demonstrated a large explicit component to the learning, evidenced by a large reduction in the reach angle when asked to remove any strategy. It could therefore be speculated that multiple mechanisms might be available when learning from binary feedback, but that if the size of the perturbation exceeds a certain magnitude an explicit strategy is required to compensate for it. Previously it has been suggested that additional learning mechanisms are recruited in response to gradually introduced visuomotor rotations when only end-point feedback is available, (Izawa and Shadmehr, 2011; Saijo and Gomi, 2010). Indeed Saijo and Gomi (2010) suggest, on the basis of an increase in reaction times, that explicit changes in motor planning occur in this paradigm. Furthermore, similarly to the results presented here, the authors also find that not all subjects are able to accomplish this. However, none of the previous studies investigating learning of rotations based on binary feedback (Izawa and Shadmehr, 2011; Pekny et al., 2015; Therrien et al., 2016) have attempted to dissect the role of implicit and explicit processes. However, learning a rotation based on binary feedback was not accompanied by a change in perceived hand position, as was found when learning was based on full visual feedback of the cursor (Izawa and Shadmehr, 2011). This could be taken as evidence that the learning described by the authors was also explicit in nature in contrast to the implicit, cerebellar-driven, adaptation.

There is increasing appreciation of the role of explicit strategies in traditional visuomotor adaptation paradigms, in which visibility of the cursor ensures that both direction and magnitude of the error are available (Bond and Taylor, 2015, 2017). The use of an ‘error-clamp’ technique has estimated the limit of implicit adaptation based on sensory prediction errors to be at around 15° (Morehead et al., 2017). Such an estimate is roughly in accordance with other estimates obtained either by the use of forcibly reduced movement preparation times (Haith et al., 2015; Leow et al., 2017), self-reporting of aiming directions (Bond and Taylor, 2015) or the difference between trials with and without strategy use (Werner et al., 2015). It is important to note in our data that all groups, with the exception of those performing the dual task, display a small amount of retention even after the removal of strategies suggesting that there is some implicit aspect to the learning. Presumably the implicit learning process triggered in the current study is distinct from the sensory prediction error driven process as here the error signal is binary in nature and provides no information about direction or magnitude of error. However, it is interesting that both implicit processes appear to be unable to compensate for rotations greater than 15-20°, with explicit strategies required for greater angles. Haith and Krakauer (2013) have proposed a theoretical framework in which model-based (strategic/explicit) and model-free (implicit) reinforcement learning processes contribute to motor learning. Our findings suggest that in the current paradigm both processes might be engaged but that the implicit process is limited in the size of rotation it can learn. It remains to be seen if this is a limitation of magnitude, as with learning from sensory prediction errors, or a limitation of speed. In other words, if the rotation was introduced more gradually or held constant for a longer period, could this implicit process account for all learning?

We measured the explicit contribution to learning via the use of an include/exclude design similar to Werner et al. (2015), which probes the contribution at the end of learning. Other approaches such as asking subjects to verbally report the aiming direction (Taylor et al., 2014) have the advantage of probing the relative contributions of implicit and explicit processes throughout learning. However, it has been suggested that this method may increase the explicit component by priming subjects that reaiming is beneficial (Leow et al., 2017; Taylor et al., 2014). Such priming may be particular powerful in paradigms like the current one as it has been shown that explicit awareness of the dimensions over which to explore is required for motor learning based on binary feedback (Manley et al., 2014). Alternatively, forcing subjects to respond at reduced reaction times can also suppresses the strategic component of adapting to a rotation (Haith et al., 2015; Leow et al., 2017). However, Leow et al. (2017) report that even at extremely short reaction times re-aiming to a single target, as used here, is still possible. In future, approaches such as measuring eye movement (Rand and Rentsch, 2016) may be beneficial to measure the explicit component during learning without priming subjects.

In order to investigate the mechanism through which subjects learnt to counter the rotation we employed the same analysis as Pekny et al., (2015). However, their study didn’t involve learning as such, as the rotation was immediately washed out. Despite this, our results are remarkably similar, in that subjects in both studies made larger and more variable changes in actions following trials in which they made an error. Sidarta et al. (2016) have also described a similar pattern of behavior when subjects attempt to find a hidden target zone based on binary feedback, with greater reductions in error following incorrect trials. Our results indicate that subjects who were unable to learn the full rotation made smaller and less variable changes in response to errors and this was primarily driven by their sensitivity to the outcome of the previous trial. Learning from errors has been suggested to be a signature of explicit reinforcement learning, in contrast to learning from success in implicit learning (Loonis et al., 2017). The finding that the difference between successful and unsuccessful subjects in the current experiments was in response to errors further supports the idea that it is the sensitivity of the explicit system that is important for this task. Interestingly, the pattern of reduced sensitivity to errors found for unsuccessful subjects in the current experiment was similar to that described for parkinsonian patients (Pekny et al., 2015). Genetic variability in various aspects of the dopaminergic system has previously been linked to differential performance in reinforcement learning (Frank et al., 2007, 2009) and the balance of model-free and model-based decision-making systems (Doll et al., 2016). Future experiments assessing if the same genetic principles apply to motor learning based on reward may be useful in not only explaining the variation in response but also cementing the links between the principles of reinforcement learning and motor learning. Interestingly, the magnitude of changes made in response to errors in a binary feedback based motor learning task was correlated with connectivity changes between motor areas, prefrontal cortex and the intraparietal sulcus (Sidarta et al., 2016). The prefrontal cortex and intraparietal sulcus have been associated with the model-based decision making system (Gläscher et al., 2010), adding further evidence for a pivotal role of explicit systems in reward-based motor learning. However, it should be noted that effects of attention and motivation cannot be ruled out in the current paradigm. Therefore, accompanying neurophysiological measures of these variables may be useful in elucidating their possible contribution.

The efficacy of the dual task paradigm employed here in preventing learning is remarkable. Dual tasks have previously been employed in conjunction with motor adaptation to visuomotor rotations (Galea et al., 2010), force-fields (Keisler and Shadmehr, 2010; Taylor and Thoroughman, 2007, 2008), as well as during the learning of motor skills (Maxwell et al., 2001) and sequence learning (Brown and Robertson, 2007). Galea et al. (2010) demonstrated that a secondary task can slow the rate of adaptation to both a gradually and abruptly introduced visuomotor rotation. Keisler and Shadmehr (2010) found that a declarative memory task could interfere with the ‘fast’ adaptation system but that a demanding cognitive task without the memory component did not. Furthermore, inhibition of the ‘fast’ process led to an increase in the ‘slow’, non-declarative process. Similarly in a sequence learning task a dual task with a declarative element increased the procedural learning suggesting that these two aspects of learning may be in competition (Brown and Robertson, 2007). It could therefore be hypothesized that the use of a dual task in the current paradigm would shift learning from explicit to the implicit system. However, the current data suggest that this did not occur and for this paradigm the explicit system is necessary to compensate for large rotations, and cannot be substituted for by an increase in the use of the implicit learning system. Whereas previous experiments have employed secondary tasks that involve more verbal systems (Galea et al., 2010; Keisler and Shadmehr, 2010; Taylor and Thoroughman, 2007), we selected the dual task which would have the maximum likelihood of disrupting the explicit system (Anguera et al., 2009; Georgopoulos and Massey, 1987). As the difficulty of the secondary task has been linked with the amount of disruption (Taylor and Thoroughman, 2008), it is also possible that the specific nature of the task may also be important and this is an interesting area for future study. One other possibility is that constant impairment of performance due to the secondary task may reduce intrinsic motivation of subjects (Liao and Masters, 2001).

The distinction between implicit and explicit reinforcement systems engaging in learning motor tasks is not merely academic. At least part of the increased interest in the addition of reward to motor adaptation and learning is due to the finding that it increases retention (Abe et al., 2011; Dayan et al., 2014, 2014; Galea et al., 2015; Shmuelof et al., 2012; Therrien et al., 2016), along with the promise this may have in a rehabilitation setting (Quattrocchi et al., 2017). However, if the benefits are primarily due to explicit or strategic processes, they may be poorly transferred to other environments and be susceptible to disruption. In line with this, it has been demonstrated that motor skills, such as golf putting or playing table tennis, are less disrupted by manipulations such as dividing cognitive load, reducing reaction times or performing in stressful situations when learnt implicitly (Liao and Masters, 2001; Maxwell et al., 2001). If the final goal of the addition of reward to motor learning tasks is to increase retention for practical rehabilitation then it may be that methods that increase the implicit contribution are required such as employing learning by analogy, reducing errors during learning or the addition of dual tasks (Liao and Masters, 2001). However, the choice and difficulty of the dual task should be made with caution as from the data presented here it may be too disruptive and ultimately prevent learning.

## Acknowledgements

The authors would like to thank Dr. Xiuli Chen for advice concerning the analysis of the data.

## Author Contributions

PH and JM designed the experiments. PH collected the data. PH and OC analyzed the data. PH, OC and JM drafted the manuscript

